# Transcriptome analysis of the binucleate ciliate *Tetrahymena thermophila* with asynchronous nuclear cell cycles

**DOI:** 10.1101/2022.07.26.501619

**Authors:** L Zhang, MD Cervantes, S Pan, J Lindsley, A Dabney, GM Kapler

## Abstract

As a prototypic ciliated protozoan, *Tetrahymena thermophila* harbors two functionally and physically distinct nuclei within a shared cytoplasm. During vegetative growth, the ‘cell cycles’ of the diploid germline micronucleus and polyploid somatic macronucleus are offset. Micronuclear S phase initiates just before cell division and is completed in daughter cells prior to the onset of macronuclear S phase. Whereas mitotic micronuclear division occurs mid-cell cycle, amitotic macronuclear division immediately precedes cytokinesis. Here we report the first RNA-seq analysis across the cell cycle of a binucleated organism. RNA was isolated at 30 min intervals across 1.5 vegetative cell cycles, starting with a macronuclear G1 population synchronized by centrifugal elutriation. Using MetaCycle, 3244 of the predicted 26,000^+^ *T. thermophila* genes were shown to be cell cycle regulated. Proteins that are required in micro- and macronuclei exhibit a single mRNA peak that correlates with their macronuclear function, while the expression of nucleus-limited protein-coding genes, including nucleoporins and importins, peak prior to their respective nucleus-specific role. Cyclin D and cyclin A/B genes showed distinct expression patterns that predict nucleus-specific functions. Clustering of periodically expressed genes revealed seven gene expression patterns. Four clusters have known PANTHER GO biological processes that are overrepresented for G1/S and G2/M phase functions. We propose that these clusters encode known and novel factors that coordinate micro- and macronuclear-specific events such as mitosis, amitosis, DNA replication and cell division.

## INTRODUCTION

Ciliates have provided a wealth of information on the structure and function of eukaryotic chromosomes, including DNA replication, telomeres and telomerase, and epigenetic regulation, including a non-coding RNA pathway that controls gene expression through programmed DNA elimination (Kataoka and Mochizuki 2011, Karrer 2012, Bracht, Fang et al. 2013). As a prototypic ciliate, *Tetrahymena thermophila*, harbors two functionally distinct nuclei-the micronucleus and the macronucleus, within the same cytoplasm. The diploid, germline micronucleus contains genetic material transmitted from parent to progeny during conjugation. It undergoes meiosis and mitosis; however, micronuclear chromosomes are never transcribed into mRNA. Post-translational histone modifications package micronuclear chromosomes into constitutive heterochromatin. The cellular phenotype is conferred by the polyploid ‘somatic’ macronucleus, where euchromatic histone modifications abound.

Upon starvation, cells of different mating type pair and produce four haploid gametic pronuclei, three of which undergo programmed nuclear death (Davis, Ward et al. 1992, Yakisich and Kapler 2004). The surviving nucleus replicates and divides, and haploid pronuclei are reciprocally exchanged between mating partners. The resulting diploid zygotic nucleus duplicates and one of the products differentiates into a macronucleus. During macronuclear development the genome is extensively reorganized (Karrer 2012). The five *T. thermophila* chromosomes undergo sequence-specific fragmentation, producing 180 non-rDNA chromosomes that are capped by telomeres and endoreplicated to ~45C. Repetitive DNA is removed by DNA breakage and re-ligation. The 21 kb ribosomal DNA (rDNA) minichromosome is amplified from 2 to 9000C. Finally, the ‘parental’ macronucleus is destroyed and progeny propagate vegetatively. Macronuclear chromosomes lack centromeres and sister chromatids randomly assort during ‘amitosis’, a poorly understood process that maintains total DNA mass and copy number within a narrow range (Doerder, Deak et al. 1992, Wong, Klionsky et al. 2000).

To accommodate the different roles of the micro- and macronuclear chromosomes, nuclear proteins must be differentially trafficked. However, shared requirements must still be met, such as chromosome packaging into nucleosomes, DNA replication and DNA repair. Micro- and macronuclear DNA replication utilize evolutionarily conserved replication machinery, including the Origin Recognition Complex (ORC), the MCM2-7 replicative helicase and other replication enzymes (reviewed in Marks, Fu and Aladjem, 2017, Mohammad, Donti et al. 2007, Donti, Datta et al. 2009). However, micro- and macronuclei are licensed to replicate at different times during the vegetative cell cycle (Allis, Colavito-Shepanski et al. 1987). Macronuclear S phase initiates prior to micronucleus S, the latter of which begins prior to cytokinesis and is completed in daughter cells. The relative timing of nuclear division is reversed-micronuclear mitosis precedes amitotic macronuclear division (Cui and Gorovsky 2006, Jacob, Lescasse et al. 2007, Cole and Sugai 2012).

Tetrahymena’s developmental and vegetative cell cycles must employ novel regulatory pathways to coordinate the fate of chromosomes in functionally distinct nuclei. Consistent with this premise, *T. thermophila* encodes 34 predicted cyclins, and 20 predicted cyclin-dependent kinases (CDKs) (Stover and Rice 2011, Yan, Dang et al. 2016, Ma, Yan et al. 2020), a subset that are only expressed in mating cells (Ma, Yan et al. 2020). To gain insight into the vegetative cell cycle of Tetrahymena, we used centrifugal elutriation to profile gene expression across 1.5 vegetative cell divisions. We report the first transcriptome analysis of the vegetative cell division cycle of a binucleate ciliated protozoan, and relate our finding to other eukaryotes, including yeast and mammals.

## RESULTS

### RNA-seq analysis across the Tetrahymena cell cycle

Centrifugal elutriation was used to generate a synchronized population of macronuclear G1 cells. This method yields >70% synchrony without chemical arrest of the cell cycle (Banfalvi 2008). Samples were collected every 30 min over 4h and RNA was isolated for high throughput sequencing. With a doubling time of ~3h, the timepoints spanned two macronuclear G1 phases (Fig. 1A). The time course was repeated twice, yielding 18 cDNA libraries generated with minimal cDNA amplification. Flow cytometry was used to assess total DNA content (Fig. 1A), EdU labeling monitored DNA replication (Fig. 1B; Fig. S1A), while DAPI fluorescence established the temporal order of micro- and macronuclear division and cytokinesis in the population (Fig. S1B). EdU labeling revealed that ~70% of macronuclei were actively replicating in the 90 min peak population (Fig. 1B, Fig. S1A) (Magiera, Gueydon et al. 2014). Micronuclear mitosis peaked at 120 min (Fig. S1B). It was immediately followed by micronuclear S phase (Fig. 1B, Fig. S1A), which was completed in daughter cells prior to the onset of the second macronuclear S phase (210 min). Of the 3216 EdU positive cells, no co-labeling of micro- and macronuclei was detected. Finally, macronuclear amitosis coincided with cytokinesis (both peaking around 180-210 min) (Fig. S1B).

**Figure 1.**
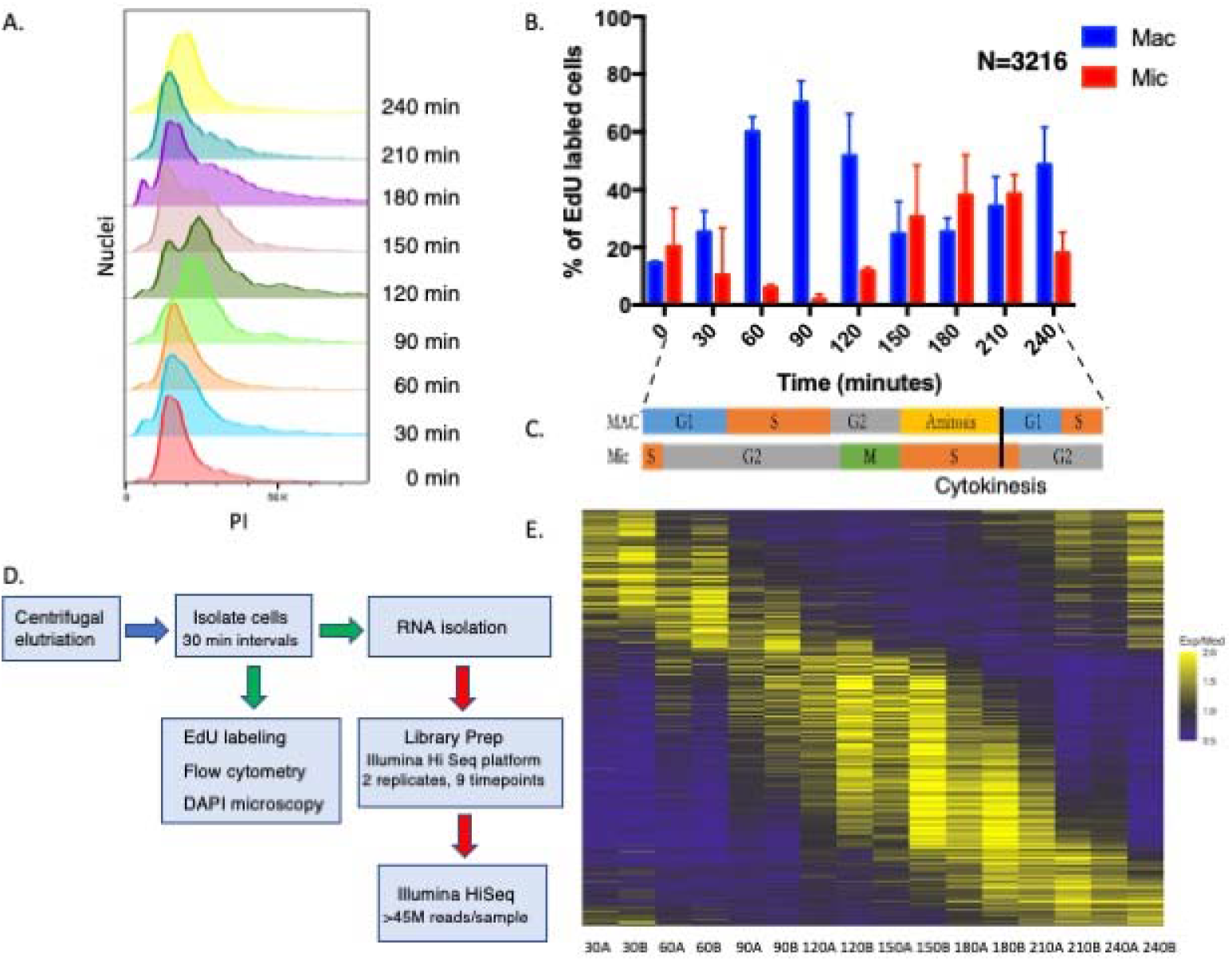
Validation of cell cycle synchrony by centrifugal elutriation. **A.** Cells isolated at 30 min intervals after elutriation were stained with propidium iodide (PI) and analyzed by flow cytometry. **B**. EdU incorporation into micro and macronuclei following centrifugal elutriation as determined by fluorescence microscopy of fixed cells. **C.** Summary time line for macronuclear and micronuclear cell cycle progression. **D.** Pipeline of sample processing for RNA-seq and cytological landmark determination. **E.** Heatmap of cell cycle regulated genes. Periodically expressed genes identified by MetaCycle for genes having 1.5-fold changes in two opposite directions.

The collective data, summarized in the Fig. 1C timeline, indicate that initiation of three of the four nuclear cycle events-micronuclear DNA replication, macronuclear DNA replication and mitosis, are temporally separated, raising the possibility that crosstalk coordinately regulates these events. In contrast, the completion of micronuclear S phase and macronuclear amitosis overlap. The macronuclear cell cycle consists of 4 phases-G1/S/G2/amitosis, while the micronuclear cell cycle has 3 phases-S/G2/M.

### Identification of cell cycle-regulated genes

To gain insight into cell cycle control of gene expression, eighteen cDNA libraries (9 per time course) were sequenced on an Illumina HiSeq 2500 resulting in ≥45 million paired end reads (43× coverage/library) with an average length of 100 bp (Fig. 1D). The paired end reads were aligned to the *T. thermophila* genome (2021 version, Tetrahymena Genome Database (http://ciliate.org/index.php/home/downloads) (Sheng, Duan et al. 2020), and the abundance of individual transcripts was computed in each sample using the HISAT2 and StringTie (Pertea, Kim et al. 2016). In all samples ≥95.5% of the reads aligned to the macronuclear genome sequence. The reads mapped to 19,946 of the 26,259 predicted *T. thermophila* genes. Quantification and statistical inference of systematic changes between timepoints were computed by DESeq2 (Love, Huber et al. 2014).

Metacycle-an N-version programming method to explore periodic data was used to identify cell cycle regulated genes (Wu, Anafi et al. 2016). Metacycle implements JTK_CYCLE (JTK) and Lomb-Scargle (LS) and integrates their results. The LS method, developed by astrophysicists as a Fourier style method for analyzing data that exhibit irregular sampling, measures the correspondence to sinusoidal curves and determines their statistical significance (Glynn, Chen et al. 2006). JTK, with its origins in statistics (Hughes, Hogenesch et al. 2010), correlates pairs of points and then computes the significance of the correlation to that of a reference curve. Because genes with subtle changes in expression are unlikely to be biologically significant, we screened for genes with a minimum of two 1.5-fold changes in opposite directions between two timepoints (p-value < 0.05) using the logarithmic fold change results by DESeq2. This analysis identified 3864 candidate genes (Fig. 1E; Supplementary Datafile 1). Using MetaCycle, 3244 of these genes were shown to exhibit periodic expression profiles with FDR <0.05. The cycling genes constitute ~11%of the total mapped genes of known or hypothetical function, and ~15% of genes expressed during the vegetative phase of the life cycle.

The results of this analysis were validated by quantitative reverse transcriptase polymerase chain reaction (qRT-PCR) for 6 genes known or anticipated to be cell cycle-regulated (Fig. S2). qRT-PCR was carried out with the same RNA preparations that were sequenced; it validated the RNA-Seq data for all examined samples. Some genes showed modest quantitative variation between biological replicates but displayed similar patterns of expression over the time course.

### Cell cycle-regulated nucleus-specific genes of known function

The specialized roles of the micro- and macronucleus are reflected in the composition of proteins that must be selectively imported into the respective nuclei. With this in mind, we looked for evidence of cell cycle regulation at the mRNA level of nucleus-specific proteins.

#### HISTONES

The nucleosomes of micro- and macronuclear chromatin share common core subunits but differ in the composition of histone variants and post-translational modifications (PTMs) that influence chromosome compaction and gene expression. The major core histones are present in both nuclei (Allis, Glover et al. 1980, Hayashi, Hayashi et al. 1984). RNA-seq profiles revealed that all core histone genes (*HH2A.1, HH2A.2, HH2B.1, HH2B.2, HH3, HH4.1* and *HH4.2*) are cell cycle regulated with a single peak in mRNA abundance during macronuclear S phase (Fig. 2A; 90 min). Variants histones H2A.Z, H3.3 and H3.4 reside exclusively in the macronucleus (Henikoff and Smith 2015). *H2A.Z* mRNA peaked during macronucleus S phase (Fig. 2B; 90 min), while H3.3 and H3.4, which serve as replacement histones after transcription-associated nucleosome removal (Yu and Gorovsky 1997, Cui, Liu et al. 2006), were constitutively expressed. Expression of the human *CenpA* homolog, *CNA1*, which functions as the micronuclear centromere-specific histone H3 variant (Cervantes, Xi et al. 2006, Cui and Gorovsky 2006) was cell cycle regulated and peaked during micronuclear mitosis (Fig. 2C; 120 min).

**Figure 2.**
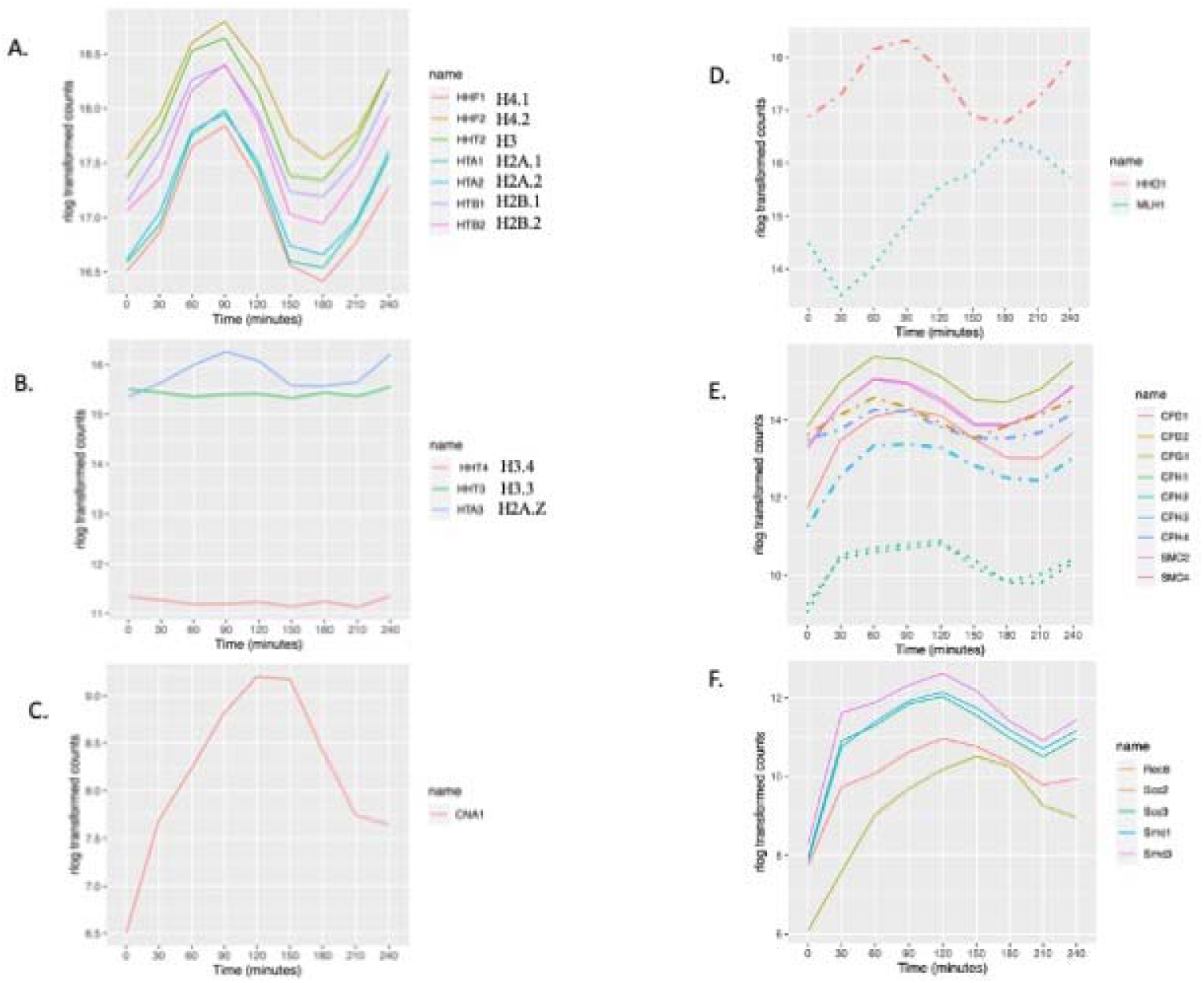
Cell cycle-regulated histone, condensin and cohesin genes. **A.** Expression profile of core histone genes. **B.** Expression profile for macronuclear-specific histone variants. **C.** Expression profile for the micronuclear-specific centromere-associated histone H3 variant, CNA1. **D.** Expression profiles for macronuclear- and micronuclear-specific H1 histones (dashed line: MAC-specific; dotted line: MIC-specific). **E.** Expression profile of cell cycle regulated condensin genes (dashed lines: macronuclear-specific; dotted lines: micronuclear-specific). **F.** Expression profile of cell cycle regulated cohesin genes (all micronuclear-specific).

The micro- and macronucleus contain different linker histones: micronuclear linker histone, *MLH1,* and macronuclear linker histone *HHO1* (Allis, Allen et al. 1984, Wu, Allis et al. 1986, Hayashi, Hayashi et al. 1987). *HHO1* is not essential for cell viability, but is critical for full chromatin compaction and maintenance of transcriptional regulation (Shen, Yu et al. 1995); (Shen and Gorovsky 1996). Whereas both linker histone mRNA levels are cell cycle regulated, they exhibited distinct expression patterns. *HHO1* mRNA peaked during macronuclear S phase (Fig. 2D; 90 min) and MLH1 peaked at the onset of micronuclear S phase (Fig. 2D; 180 min).

#### HISTONE MODIFYING ENZYMES

Micro- and macronuclear histones are subjected to different PTMs (Vavra, Allis et al. 1982). The micronucleus is enriched for H3K27me3 and completely lacks the euchromatic mark H3K4me3 (Liu, Taverna et al. 2007, Taverna, Ueberheide et al. 2007). The macronucleus contains both H3K27me3 and H3K4me3 (Papazyan, Voronina et al. 2014). During vegetative growth, histone H3 acetylation (Chalker, Meyer et al. 2013) and deacetylation (Wiley, Myers et al. 2005), and Txr1p-catalyzed H3K27 monomethylation occur in the macronucleus (Gao, Xiong et al. 2013), whereas H3K23 trimethylation is micronucleus-specific (Papazyan, Voronina et al. 2014).

Five known histone modifying enzyme genes are cell cycle regulated. mRNA levels for the two histone acetyltransferases, *HAT1, HAT2,* associated with transcription activation, peaked in early macronuclear S phase (Fig. 3A, 60 min). Expression of macronuclear-specific *TXR1*, peaked sharply during macronuclear G1 phase (Fig. 3A, 30 min). Two of the three Drosophila Enhancer of Zeste (EZ) are also cell cycle regulated. The histone methyltransferase genes *EZL1, EZL2* and *EZL3* are responsible for H3K27 di- and tri-methylation. Whereas TXR1p is macronuclear-limited, EZL2p and EZL3p are not. EZL2p mediates di- and trimethylation of H3K27 in early conjugants and vegetatively growing cells (Gao, Xiong et al. 2013, Zhang, Molascon et al. 2013). It is required for global transcriptional silencing in the micronucleus during vegetative growth, and localized repression of gene expression in the macronucleus. The role of EZL3p is less well-defined. *EZL2* and *EZL3* mRNA levels are cell cycle-regulated; they produced broad peaks that spanned late micronuclear S phase and macronuclear S phase (Fig. 3A, 30-120 min). Consistent with the role of EZL1p in heterochromatin formation and DNA elimination in the developing macronucleus (Liu, Taverna et al. 2007, Xu, Zhao et al. 2021); EZL1 expression was low in vegetative cells and did not cycle.

**Figure 3.**
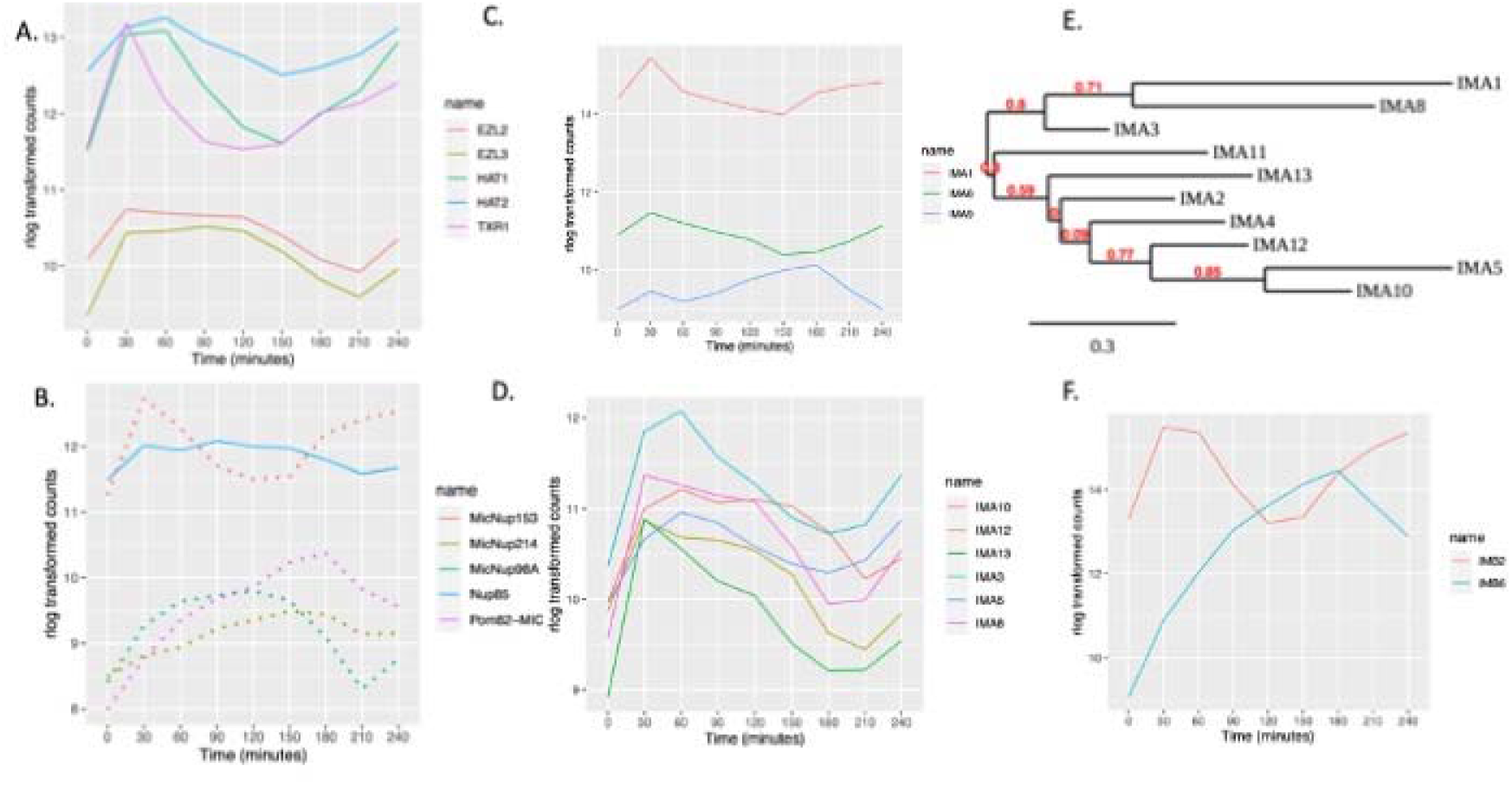
Expression profile of histone modifiers, and macro- or micronuclear-specific nucleoporin genes. **A.** Cell cycle-regulated histone modifiers (see text for gene names). **B.** Cell cycle regulated nucleoporins (dotted lines: micronuclear-specific, solid lines-present in both nuclei). **C.** Cell cycle regulated expression of macronuclear of non-specific importin-alpha genes; IMA1 (macronuclear-specific), IMA6 and IMA9. **D.** Cell cycle regulated expression of micronuclear importin-alpha genes. **E.** Phylogenetic tree analysis of importin alpha genes. **F.** Cell cycle regulated importin beta-like proteins.

Histone H3 lysine 4 (H3K4) methylation is mainly catalyzed by MLL1p and is associated with transcription activation (Cao, Chen et al. 2010). *MLL1* is not periodically expressed. Similarly, genes responsible for macronuclear histone H3 demethylation (*THD1, SIN3/TTHERM_00450950, RXT2/TTHERM_00992830, SAP30, PHO23/TTHERM_000046389, SAP18/TTHERM_00469050*) and histone H2 mono-ubiquitinylation (*RIN1/TTHERM_00263030;* UB3 ligase) are not cell cycle regulated (Zhang, Gao et al. 2014, Nabeel-Shah, Garg et al. 2021).

#### CONDENSINS AND COHESINS

The Structural Maintenance of Chromosomes (SMC) complex plays a central role in chromosome segregation. Its subunits and nuclear localization have been well studied in *Tetrahymena* (Uhlmann 2016). *Tetrahymena* has a reduced set of SMC complexes, containing a single cohesion that resembles the meiotic cohesin of other eukaryotes, and two heterodimeric condensin I complexes that are related to condensins that mediate metaphase compaction in yeast and higher eukaryotic chromosomes ((Howard-Till, Tian et al. 2019). *T. thermophila* condensin I complexes are targeted to either the micro-or macronucleus and are distinguished by their kleisin/Gph subunit. Micronuclear condensins contain CPH1p and CPH2p, whereas macronuclear condensins contain CPH3p and CPH4p. CPH5p localizes to the macronucleus during development. All condensin genes are cell cycle regulated except CPH5p, which has a critical (possibly primary) role in mating cells. Consistent with previous research (Howard-Till, Tian et al. 2019), mRNA levels for the two micronuclear kleisin genes, *CPH1* and *CPH2*, peak at 120 min, corresponding to mitosis, and their expression patterns differ from the macronuclear condensin subunits (Fig. 2E). Furthermore, genes encoding subunits that are present in both the micro- and macronucleus were cell cycle regulated (Fig. 2E; CPD1, CPD2 and CPG3, SMC2 and SMC4).

*Tetrahymena* has a minimal cohesin complex consisting of SMC1p, SMC3p, SCC3p, and REC8p, that localizes specifically to the micronucleus (Ali, Loidl et al. 2018). This complex performs all necessary functions for mitosis and meiosis. SCC2p, which in other organisms is part of a heterodimeric complex (Scc2p/Scc4p) that helps load cohesin onto chromatin, is not required for chromosomal association of cohesin, but *Tetrahymena* Rec8p is hypo-phosphorylated in its absence (Ali, Loidl et al. 2018). Consistent with the random segregation of sister chromatids in the amitotic macronucleus, cohesins are not targeted to the macronucleus. In our dataset, peak gene expression of all the cohesin subunits occurs during micronuclear mitosis. Cohesin subunit gene expression patterns cluster closely together (Fig. 2F, 120 min), except for *SCC2* whose mRNA level peaked slightly later (Fig. 2F, 150 min).

#### NUCLEAR PORE COMPONENTS

Gene duplication and sub-functionalization has generated 4 micronuclear-specific, 5 macronuclear-specific, and 1 shared nuclear pore or nuclear transmembrane protein (MicNup98Ap, MicNup153p, MicNup214p, MicPom82p; MacNup98Ap, MacNup98Bp, MacNup153p, MacNup214p, MacPom121p; shared Nup85p). Minimally, there are 16 additional shared protein components of micro- and macronuclear pores (Iwamoto, Mori et al. 2009) (Iwamoto, Osakada et al. 2017). Iwamoto et. al. proposed that nuclear-restricted subunits confer selective cargo transport. Our data revealed that the micronucleus-specific nucleoporin genes (*MicNup98A, MicNup214* and *MicNup153*) and the micronuclear transmembrane protein, *MicPom82,* are cell cycle regulated (Fig. 3B). *MicNup153* expression peaked at micronucleus S/G2 phase and *MicPom82* and *MicNup214* peaked at micronuclear S phase. *MicNup98A* peaked at micronucleus mitosis. In contrast, none of the macronuclear-specific or shared pore protein subunit or transmembrane protein genes were cell cycle regulated.

#### IMPORTINS AND EXPORTINS

Trafficking of proteins into the nucleus is mediated by the interaction between nuclear localization signals (NLSs) in cargo proteins and chaperones, termed importins (reviewed in Wing, Fung et al. 2022). Importins alpha and beta form heterodimers to transport cargoes through the nuclear pore. Tetrahymena encodes 10 importin alphas, 1 that localizes to the macronucleus (IMA1) and 9 that localize to the micronucleus (Malone, Falkowska et al. 2008). In addition, there are 2 importin-related proteins, IMA6p and IMA9p of unknown function. Macronuclear-associated *IMA1* is expressed at much higher levels than its micronuclear counterparts but did not meet the criteria for cell cycle regulation (Fig. 3C). In contrast, 6 of the micronuclear importin genes, *IMA3, IMA5, IMA8, IMA10, IMA12* and *IMA13,*are cyclically expressed. Their mRNAs exhibit different profiles that in general peak in the 30-90 min window (Fig. 3D). *IMA10,* with peak expression at the micronuclear G2/M transition, is an essential gene that is required for chromosome segregation during mitosis and meiosis (Malone et al., 2008). Whereas *IMA2* and *IMA11* expression profiles met the 1.5-fold cyclic change in abundance, they were not detected by MetaCycle.

Phylogenetic tree analysis supports two importin-alpha gene family groupings: *IMA1, IMA2, IMA3, IMA8, IMA11* and *IMA13* form one group. Our data shows that their cell cycle regulated genes all peak during the micronuclear S/G2 window. *IMA4, IMA5, IMA10 and IMA12* form the second group, and their expression peaks at micronuclear G2/M (Fig. 3E) (Dereeper, Guignon et al. 2008). The different importin-alpha expression profiles, their phylogenetic relationships, and functional data suggest that these gene family members serve non-redundant roles in protein trafficking. Finally, importin-alpha related genes of unknown function, *IMA6* and *IMA9,* are both cell cycle regulated (Fig. 3C).

*T. thermophila* encodes 11 importin-beta proteins, none of which are nucleus-specific (Malone et al., 2008). IMB4 and IMB8 have a pronounced macronuclear bias. IMB2 protein accumulates in the cytoplasm in a GFP-tagged overexpression strain, while IMB6 localizes to both the micro- and macronucleus (Malone, Falkowska et al. 2008). Only *IMB2* and *IMB6* were cell cycle regulated (Fig. 3F).

### Identification of potential nucleus-specific cell cycle regulatory proteins

Major drivers of eukaryotic cell cycle progression include cyclins, cyclin-dependent kinases (CKDs) and E2F transcription factors. These genes are typically regulated at multiple levels, including transcription (Harashima, Dissmeyer et al. 2013, Malumbres 2014). To identify potential micro- and macronucleus-specific cell cycle regulators, we correlated the expression profiles of predicted cyclin, CDK and E2F genes to known functions of their yeast and mammalian homologues.

Tetrahymena encodes at least 34 predicted cyclin genes (Stover and Rice 2011, Yan, Dang et al. 2016). Phylogenetic analysis suggests that 11 are cyclin D orthologues and 10 are members of the cyclin A/B family (Stover and Rice 2011). Cyclin D regulates the G1/S transition (Grana and Reddy, 1995). Our RNA-seq analysis revealed cell cycle-regulation of 4 cyclin D transcripts. *CYC7, CYC12,* and *CYC22* mRNAs peaked just prior to the macronuclear G1/S transition (Fig. 4A, 30 min), while *CYC14* mRNA was most abundant prior to the onset of micronuclear S phase (Fig. 4A, 120 min). *CYC3* and *CYC26* did not exhibit a 1.5-fold change in opposite directions at two timepoints, but met the Metacycle criteria for oscillating genes, peaking at 180 min (micronuclear S phase). *CYC9* and *CYC2* mRNA levels were low and did not oscillate in vegetative growing cells, consistent with previously defined roles during development (Yan, Dang et al. 2016, Ma, Yan et al. 2020) (Suppl. Fig. S3A). The remaining cyclin D family members, *CYC4, CYC13* and *CYC25,* were constitutively expressed during vegetative growth.

**Figure 4.**
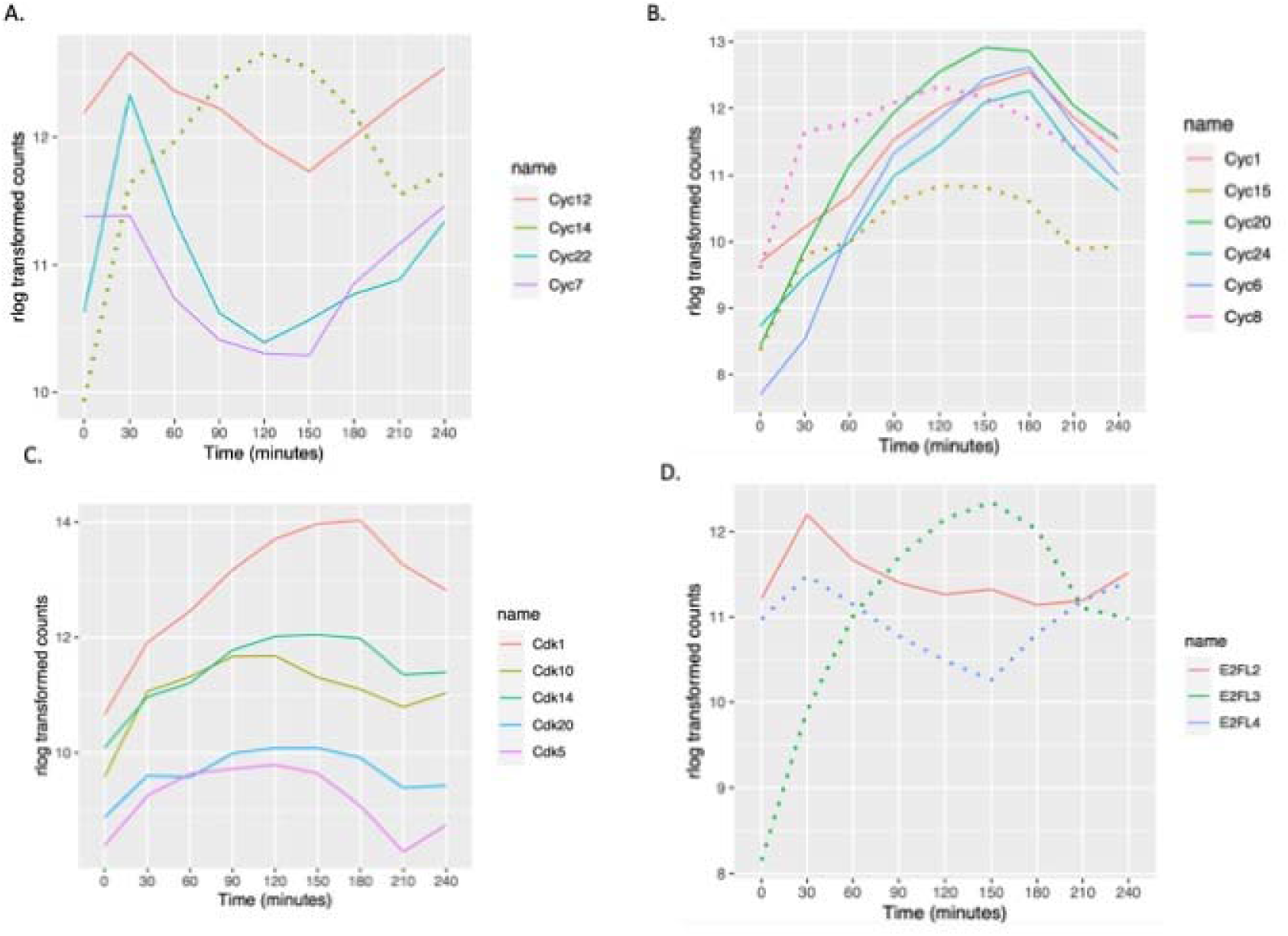
Oscillating cyclin, CDK and E2F mRNA abundance profiles. **A.** Cell cycle regulated expression of predicted cyclin D gene family members. **B.** Cell cycle regulated expression of predicted cyclin A/B genes. Solid lines: pattern 1 of cyclin A/B; dotted lines: pattern 2 of cyclin A/B. **C.** Cell cycle-regulated CDK genes. **D.** Cell cycle-regulated E2Fs. Solid line: potential E2F activator; dotted lines: potential E2F repressors and atypical E2F with two DNA-binding domains.

Cyclin A/B family members regulate the S/G2 or G2/M transitions (Grana and Reddy 1995). Predicted Tetrahymena cyclin A/B gene family members exhibited two distinct oscillating patterns. mRNA levels for *CYC1, CYC6, CYC20, CYC24* peaked during macronuclear amitosis and cytokinesis, when the micronucleus is in S phase (Fig. 4B, 150 and 180 min; Fig. S3B). A similar oscillation was observed for CYC18 by Metacycle, but the 1.5-fold change threshold was not met. In contrast, the *CYC15* and *CYC8* (and possibly *CYC10*) peaks coincided with micronuclear mitosis (Fig. S3B, 120 min). *CYC11* was not cell cycle-regulated, although its expression level was comparable to other A/B family members. As expected from previous functional studies (Yan, Dang et al. 2016), the meiosis-specific A/B cyclin gene, *CYC17,* was constitutively expressed at very low levels during vegetative growth.

Twenty-two putative CDK/CDK-like genes have been identified in the *T. thermophila* (Ma, Yan et al. 2020). Five of these genes are cell cycle-regulated at the mRNA level. CDK1 mRNA was highest at 180 min (Fig. 4C, late micronuclear S phase/macronuclear division/cytokinesis). *CDK5, CDK10, CDK14, CDK20* exhibited similar broad oscillating peaks (Fig. 4C), and the remaining CDKs were constitutively expressed (Fig. S3C). *CDK19* expression was very low in vegetative cells, consistent with its conjugation-specific role (Ma, Yan et al. 2020).

Gene predictions identified 7 putative *T. thermophila* E2F gene family members (Zhang, Tian et al. 2017). E2F transcriptional activators drive expression of S phase-specific genes, increasing in G1 and peaking during S phase. E2F repressors have the opposite effect and peak during G2 phase (Thurlings and de Bruin 2016). Tetrahymena *E2FL1* and E*2FL2* have conserved E2F/DP family winged-helix DNA-binding and CC-MB domains, features of transcriptional activators. *E2FL3* and *E2FL4* are most closely related to the mammalian repressors, *E2F7* and *E2F8*. *E2FL1* expression was constitutively low in vegetative growing cells, consistent with its role in meiosis (Fig. S3D) (Zhang, Tian et al. 2017). *E2FL2* mRNA peaked at macronuclear G1 phase, suggesting that it is involved in activation of S phase-specific genes (Fig. 4D, 30 min). The putative repressor gene, *E2FL3* also peaked at 30 min, while *E2FL4* peaked at 150 min corresponding to exit from macronuclear S phase. *DPL1, DPL2* and *DPL3* did not meet the criteria for cell cycle regulation, however, their expression profiles tracked with *E2FL2* (Fig. S4D).

In mammalian cells, MuvB complexes interact with E2F4/5, DP1/2 and an RB-like protein to generate transcriptional repressor complexes that inhibit expression of cell cycle regulated genes (Litovchick, Sadasivam et al DeCaprio (2007) Mol Cell; Sadasivam and DeCaprio, 2013, Nat Rev Cancer). In contrast, to the predicted Tetrahymena E2F repressors (*E2FL3* and *E2FL4*), none of the *T. thermophila* paralogues and Reb1 (Rb-like)-associated proteins identified by a proteomic-based approach (Nabeel-Shah, Garg et al. 2021) are cell cycle regulated at the level of transcription (RebL1, Lin9, Lin54, Anqa1, Jinn1, Jinn2).

### Periodically expressed gene clusters

Using the noise-robust soft clustering algorithm (Futschik and Carlisle 2005) periodically expressed genes could be parsed into 7 clusters, of which 4 have known PANTHER GO biological process overrepresentation (Fig. 5A). The other 3 clusters have unknown biological functions. The later result is not unexpected, since only 57% of the cell cycle-regulated *T. thermophila* genes have assigned GO biological processes (1742 out of 3032). mRNA abundance for genes in cluster 7 was maximal during macronuclear G1 and S phase. Cluster 5 genes were maximally expressed during macronuclear S and G2 phase. Genes in cluster 1 were maximally expressed during macronuclear G2 and amitosis. Genes in cluster 2 were maximally expressed during amitosis and cytokinesis.

**Figure 5.**
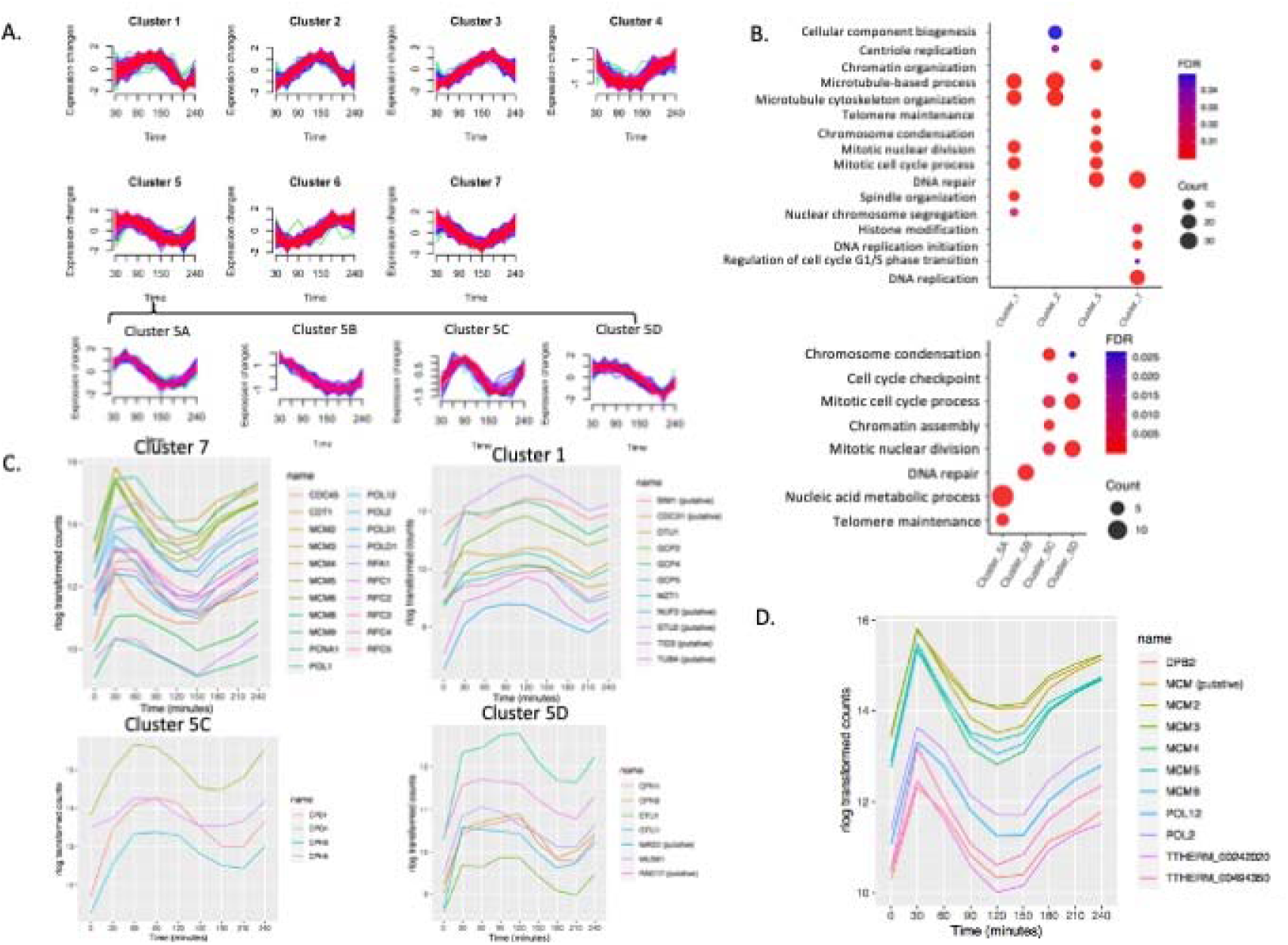
Cluster and GO enrichment analysis. **A.** Clusters of 7 distinct periodically gene expression profiles and 4 sub-clusters. **B.** Overrepresented GO biological processes of four clusters. Overrepresented GO biological processes of sub-clusters of cluster 5. **C.** Gene expression profiles DNA replication protein-coding genes within cluster 7. Genes expression profile of mitotic genes within cluster 1. Gene expression profile of mitotic genes of cluster 5C. Gene expression profile of mitotic and cell cycle checkpoint genes of cluster 5D. **D.** Co-regulation analysis of MCM6 mRNA abundance: top 10 co-regulated genes.

#### Cluster 7

GO overrepresentation within the 531 gene cluster 7 (macronuclear G1/S, late micronuclear S) showed a highly significant enrichment for processes linked with S phase, including DNA replication (P=1.00E-20), DNA repair (P=1.03E-18), regulation of the G1/S transition (P=0.04) and histone modifiers (P=3.36E-3) (Fig. 5B; Supplementary Datafile 2). Genes in cluster 7 include several factors involved in DNA replication, such as various polymerases (*POL1, POL12, POL2, POL31, POLD1*), proliferating cell nuclear antigen (*PCNA1*)and other proteins necessary to initiate DNA synthesis (*MCM2-6, CDT1,* DNA Replication Factor 1 (*RFA1*), *CDC45,* and *RFC1-5*) (Fig. 5C). DNA repair and DNA damage recognition factors known to cooperate in DNA replication were also identified, including RAD complex components (*RAD50*, *RAD51*, *RAD54*), *MCM8*, *MCM9*, *BRCA1* and *BRCT2*. Importantly, genes that regulate or drive the G1/S transition, including cyclin Ds (*CYC12*, *CYC22*), *E2FL2* and *E2FL4,* reside in cluster 7. *CDC6* and *ORC1* displayed two 1.5-fold changes at two opposite directions but failed to be detected by MetaCycle; their expression level peaked at 30 min but did not have display a clear peak at the second G1 phase. This may be due to the lack of synchrony at later time points (Fig. S5). It is worth noting that a putative a cell cycle checkpoint serine-threonine kinase-*DUN1* homolog (*TTHERM_000851660*) which is required for transient G2/M arrest after DNA damage, *CDC14,* which is required for mitotic exit, and the non-SMC mitotic condensation complex subunit, *CPD2*, are also in cluster 7. Mitosis-related genes in this cluster may function in the G2/M phase transition of the germline micronucleus.

#### Cluster 5

284 genes are represented in Cluster 5, whose expression peaks during macronuclear G2/micronuclear mitosis, with GO association enrichments that include the as mitotic cell cycle process (P=1.96E-07), mitotic nuclear division (P=1.90E-07, chromosome segregation (P=1.56E-08), chromosome condensation (P= 2.37E-07), telomere maintenance (P=5.32E-05), DNA repair (P=2.03E-07) and chromatin organization (P=8.90E-06) (Fig. 5B, upper panel). Sub-clustering of cluster 5 allowed more specific cell cycle phase associations (Fig. 5B, lower panel; Supplementary Datafile 3).

Genes in cluster 5A peak at early MAC S phase (60 min) and are enriched for biological processes such as telomere maintenance (P=2.37E-06) and nucleic acid metabolic process (P=7.94E-07) (Supplementary Datafile 4). Cluster 5B consist of genes whose expression peaks during macronuclear G1 phase (30 min). The associated GO term is DNA repair (P=5.32E-07) (Supplementary Datafile 5).

Cluster 5C consists of genes that are up-regulated at late macronuclear S phase/micronuclear mitosis (90 min), and whose GO associations are enriched for the terms chromosome condensation (P=7.44E-07), chromatin organization (P=2.32E-06), mitotic cell cycle process (P=3.68E-03) and mitotic nuclear division (P=3.59E-03) (Fig. 5B; Supplementary Datafile 6). The two macronucleus-specific condensin genes (CPH3 and CPH4) and other two subunits of the condensin complex (*CPD1, CPG1*) (Till, Tian et al. 2019) are in cluster 5C (Fig. 5C). The PANTHER annotated pathway of mitotic cell cycle process and mitotic nuclear division are due to those four genes. Cluster 5C also contains several histone genes in the chromatin organization pathway (*HHF2, HTA1, HTA2, HTA3, HTB1, HTB2*).

Cluster 5D contains genes with high expression levels through micronuclear G2 phase and mitosis, that downregulated at amitosis/cytokinesis. This cluster is enriched for biological pathways such as cell cycle checkpoint (P=4.49E-03), chromosome condensation (P=2.64E-02), mitotic nuclear division (P=2.31E-04) and mitotic cell cycle process (P=3.47E-04) (Fig. 5B, lower panel; Supplementary Datafile 7). The two micronuclear-specific condensin genes (CPH1 and CPH2) are in cluster 5D. Cell cycle checkpoint genes in cluster 5D include *MUS81/RAD17 (TTHERM_000191179*) and *MAD3 (TTHERM_00393260*), a subunit of the spindle-assembly checkpoint complex (Fig. 5C). Other genes in chromosome segregation include epsilon-tubulin (*ETU1*) and gamma-tubulin (*GTU1*). Epsilon-tubulin localizes to the pericentriolar material and is required for centriole duplication and microtubule organization. Its recruitment to the new centriole can only occur after exit from S phase (Chang and Stearns 2000, Chang, Giddings et al. 2003).

#### Cluster 1

The 342 genes in cluster 1 (micronuclear mitosis, macronuclear G2) were highly enriched for mitosis-related biological processes such as mitotic cell cycle process (P=2.95E-06), mitotic nuclear division (P=1.48E-06), spindle organization (P = 2.96E-05) and microtubule-based processes (P=2.98E-06). The mitosis-related biological process GO term-chromosome segregation has a higher P value (P=2.25E-02), partly because several cohesin genes are not annotated in PANTHER (SMC1, SMC3, SCC3 and REC8), (Fig. 5B; Supplementary Datafile 8). Spindle organization includes the mitotic-spindle organizing protein (Mzt1), putative kinetochore protein (Nuf2p), putative Microtubule-associated protein (Stu2p), putative microtubule-binding proteins (Bim1) and hypothetical protein (TTHERM_00493000); and several genes encoding the gamma-tubulin complex (*GCP2, GCP4, GCP5*) (Fig. 5C). Predicted mitotic genes, such as *CDC31, TUB4,* and *TID* are also in cluster 1. Delta-Tubulin 1 (*DTU1*) which is found in association with the centrioles and associating with only the older of the centrosomes in a newly duplicated pair is also in this cluster (Chang and Stearns 2000). This group includes Cyclin A/B (*CYC8*) and Cyclin D (*CYC14*) (Fig. 4A; Fig. 4B) and all genes encoding the minimal cohesin complex components (*SMC1, SMC3, SCC3* and *REC8*) (Fig. 2F).

#### Cluster 2

This group contains 564 genes whose expression peaked at 150 min (macronuclear amitosis/cytokinesis). GO terms are associated with microtubule-based process (P=6.42E-08), cytoskeleton organization (P=1.94E-06), cellular component biogenesis (P=4.9E-02) and centriole duplication (P=3.33E-02) (Fig. 5B; Supplementary datafile 9). Three genes in centriole duplication from PANTHER are *CPAP1, TTHERM_00532450* (centrosome protein putative), and *TTHERM_01113110* (DHHC zinc finger protein). Sister Chromatid Cohesion protein 2 (SCC2), the tubulin ligases, *TTLL1, TTLL2, TTL10,* components of the anaphase-promoting complex (*APC1, APC2, APC10*), and two cell cycle regulators, *CYC15* and *E2FL3,* are in this cluster. These data suggest a degree of functional conservation between mitosis and amitosis.

### Co-expression profiling analysis identifies functionally regulated genes

Genes with similar expression patterns during the cell cycle frequently participate in the same pathway. Weighted gene co-expression network analysis (WGCNA) is a systems biology method for describing the correlation patterns among genes across RNA-seq samples (Zhang and Horvath 2005, Langfelder and Horvath 2008). Co-expression networks have been found useful for describing the pairwise relationships among genes. We identified the top 10 co-expression profiles for each cell cycle-regulated gene using WGCNA (see Supplementary Data File 10). The results can be used for data exploration or gene screening of a given pathway. To illustrate the utility of co-expression, we used MCM6 as our query. This gene is co-regulated with other genes involved in the initiation of DNA replication and replication fork progression in other eukaryotes (Cho, Campbell et al. 1998, Menges, Hennig et al. 2003, Grant, Brooks et al. 2013) (Fig. S4). This survey identified other subunits of the MCM complex (*MCM2, MCM3, MCM4, MCM5, MCM6*) and the replication polymerase (*POL2, POL12, DPB2*). It also identified a putative MCM gene family member that was not previously annotated (*TTHERM_000011759*) (Fig. 5D). The two remaining ‘top 10’ co-regulated genes have unknown functions, one of which contains a serine/threonine kinase domain. Strong co-expression correlations were observed in particular for genes involved in other aspects of DNA metabolism as well as nucleocytoplasmic scaffolding proteins.

## DISCUSSION

As an early branch in the eukaryotic lineage, ciliated protozoa have evolved a truly unique mechanism for partitioning chromosomal functions through the genesis of two organelles with non-overlapping roles-the micronucleus and macronucleus. The only known function of the micronucleus is transmission of chromosomes from parent to progeny during conjugation. In sharp contrast, the macronucleus is sole dedicated to gene expression. Development of a new macronucleus involves massive reorganization of the micronuclear chromosomes, including programmed elimination of repetitive and unique DNA sequences using an RNA-guided mechanism and genome-wide reprogramming of heterochromatin into euchromatin (Yao and Chao 2005). Once macronuclear development is complete, the parental macronucleus undergoes autophagic programmed nuclear death (Liang, Xu et al. 2019) and references therein).

One consequence of the separation of micro- and macronuclear functions is the creation of a complex vegetative cell cycle in which the two nuclei replicate and partition their respective chromosomes at different times. Using centrifugal elutriation, RNA-Seq profiling, EdU labeling and DAPI analysis, we have expanded our understanding of the relationship between the two nuclear cycles. Our molecular and cytological data support a model for coordinated signaling between the micro- and macronucleus. Differences in the temporal expression of cyclin D family members argue for the separate licensing of two discrete S phases, in which the macronucleus (pure euchromatin) initiates DNA replication first and the micronucleus (pure heterochromatin) follows once macronuclear S phase is complete. Crosstalk between nuclei is supported by our observation that none of the 3000+ EdU-positive cells that we scored incorporated label into both the micro- and macronucleus. Whereas the macronucleus replicates first, the micronucleus undergoes mitosis prior to amitotic macronuclear division. Following mitosis, the micronucleus immediately enters S phase, which is completed in daughter cells. Amitosis of the macronucleus coincides with cytokinesis (Fig. 1C).

Our RNA-Seq analysis of the *T. thermophila* vegetative cell cycle is the first such analysis in a binucleated species. In total, steady state mRNA levels of over 3000 genes were shown to be under cell cycle control. 16% of vegetative expressed genes exhibited cyclic oscillations that peaked at different stages of the cell cycle. This percentage is somewhat higher than S*. cerevisiae,* and cultured plant (*Arabidopsis thaliana*) and human cells analyzed by microarray (range 5-10%). This repository of information can be used to identify genes that control fundamental processes that occur in both the micro- and macronucleus but may be differentially regulated or mediated by different proteins. One such example is mitotic and amitotic chromosome transmission. Since many mitotic genes in yeast, higher eukaryotes and Tetrahymena are cell cycle-regulated (Cho, Campbell et al. 1998, Spellman, Sherlock et al. 1998, Rustici, Mata et al. 2004, Giotti, Joshi et al. 2017), it is possible that the same would hold for amitosis. Proteomic studies previously demonstrated that two of the four condensins localize to the macronucleus (Fig. 2E; *CPH3, CPH4*). In stark contrast, none of the cohesins are targeted to the amitotic macronucleus (Howard-Till, Tian et al. 2019). All cohesin subunit genes and micronuclear condensins (*CPH1, CPH2*) are cell cycle-regulated, peaking prior to micronuclear mitosis (Fig. 2E, Fig. 2F). Meanwhile, cell cycle-regulated expression of macronuclear condensing genes is maximal just before amitosis (Fig. 2D). The identification of co-regulated genes can provide an avenue to pursue the amitotic program further. Furthermore, the identification of known chromosome segregation genes with peak expression after micronuclear mitosis may provide a path forward towards gaining insights into the amitotic program.

Cyclin-CDK complexes are the master regulators of the eukaryotic cell cycle (Grana and Reddy 1995). Tetrahymena encodes 34 putative cyclin genes, a subset that are exclusively expressed during conjugation (Stover and Rice 2011, Xu, Wang et al. 2016, Yan, Dang et al. 2016, Ma, Yan et al. 2020). Of the 11 predicted cyclin D genes, *CYC7, CYC12* and *CYC22* expression peaks at the macronuclear G1/S transition, and CYC14 peaks prior to the onset of micronuclear S phase. In keeping with the known role of cyclin D, our data suggests that these paralogs regulate entry into macro- and micronuclear S phases, respectively. Cyclin A/B family members regulate the S/G2 and G2/M transitions. Two A/B family members peak prior to mitotic micronuclear division, *CYC15, CYC8* (and possibly *CYC10*). Four cyclin A/B genes (*CYC1*, *CYC6*, *CYC20*, *CYC24*) peak later in the cell cycle suggesting they may regulate macronuclear amitosis. The non-essential vegetative role of the micronucleus and phenotypic assortment of alleles in the amitotic polyploid macronucleus (Turkewitz, Orias et al. 2002) provide opportunities to determine the functional roles of vegetative cyclins using reverse genetic approaches, similar to prior work on ORC and the ATR checkpoint kinase (Smith, Yakisich et al. 2004, Yakisich, Sandoval et al. 2006, Lee, Meng et al. 2015).

The diploid (2C) heterochromatic micronucleus contains 5 chromosomes, ~1/3 of which consists of repetitive DNA (mostly transposable elements), and non-redundant sequences, all of which are excised from chromosomes during macronuclear differentiation (Hamilton, Kapusta et al. 2016). The 181 euchromatic macronuclear chromosomes endoreplicate and are subsequently maintained at ~45C. Whereas core histone proteins (H2A, H2B, H3, H4) are present in micro- and macronuclei, there is a single increase in cell cycle-regulated gene expression that peaks prior to macronuclear S phase, when the demand for these proteins is highest (Fig. 2A-D). The same expression pattern is observed for pre-RC components and the core DNA replication machinery (Fig. 5A; Fig. S5). Since the DNA mass in the macronucleus is ~20-fold greater than the micronucleus (45C vs. 2C), the need for two waves of gene expression seems unnecessary. While core histones could be imported into both nuclei concurrently, a more efficient approach would be to regulate the timing and amount of core protein import into micro- and macronuclei. We speculate that the cell cycle-regulated transcription of nuclear pore subunits, importins and cyclin/CDK complexes (Fig. 3; Fig. 4) in conjunction with post-translational modification of these and other factors regulate protein trafficking into micro- and macronuclei. To the best of our knowledge, our work provides the first example for cell cycle-regulated expression of genes encoding components of the nuclear pore and cargo transporter proteins.

In contrast to core histone subunits, centromeric histone H3 variant, *CNA1,* micro- and macronuclear-specific H1 linker histones (MLH1 and HHO1, respectively) the macronuclear H2A.Z histone variant, and macronucleus-specific chromatin modifiers (*TXR1*, *HAT1, HAT2, EZL2* and *EZL3*) peak with the onset of their respective nuclear S phase. This is consistent with temporal linkage between demand and supply.

A powerful application of transcriptomics is the establishment of interaction or functional networks. We employed two strategies for our analysis of cell cycle-regulated genes. The first approach involved noise-robust soft clustering analysis-an unsupervised learning technique designed to reveal structures hidden in large time-course gene expression data sets. The second approach-weighted gene co-expression network analysis (WGCNA), uses a different algorithm to find clusters by describing the correlation patterns between genes. The cluster approach identified 7 expression patterns that account for the 3032 cell cycle-regulated genes identified by MetaCycle. Due to the relatively poor annotation of the predicted 26,259 genes in this early branching eukaryote, only four of the seven clusters were populated with genes with assigned GO terms. Of note-there was significant overlap for GO terms in clusters 1 and 2 for microtubule-based processes that occur at different stages of the cell cycle, suggesting that mitosis and amitosis utilize related machinery to transmit chromosomes to daughter nuclei. Unexpectedly, GO terms for centriole replication and cellular component biogenesis were enriched at the end of the cell cycle when macronuclear amitosis occurs. Delving deeper into the clustered genes through WGCNA co-expression to study genes of unknown function could provide new insights into mechanisms that are conserved throughout evolution, as well as novel mechanisms that have evolved to solve species-specific problems, such as the maintenance of genome balance in the absence of mitosis, the temporal licensing of DNA replication in binucleate species, and the epigenetic control of replication initiation in euchromatic and heterochromatic chromosomes with nearly identical unique sequence composition.

## METHODS

### Synchronization by centrifugal elutriation

CU428 cells (1.5L) were grown to ~1×10^5^ cells/ml in 2% PPYS/1000U/ml pen-strep (Sigma Aldrich) at 30°C. Elutriation was used to obtain a synchronized G1 cell population as previously described (Liu et al. Frontiers Cell Dev Biol 2021). Cells were subsequently collected at 30 min intervals spanning 1.5 cell cycle, 240 min). For each timepoint, 20 ml of cells were collected; 5 ml for flow cytometry and DAPI analysis, and 15 ml for RNA isolation. Centrifuged cell pellets were washed in 10 mM Tris pH 7.4 prior to flow cytometry and RNA isolation. Two biological replicates were used to generate 18 cDNA libraries.

### Flow cytometry and EdU labeling

Cells were prepared for flow cytometry as previously described (Sandoval et al.). CU428 cells were synchronized using centrifugal elutriation as described above. 10 min before each time point, 2 ml of cells were labeled for 20 min with 25 uM 5-ethylene-2’-deoxyuridine (EdU) (Life Technologies), washed twice with PBS, fixed with 4% PFA for 5 min, rinsed twice with PBS, and stored at 4°C. To visualize EdU incorporation, 100 ul samples were spotted onto coverslips, and then let dry for 2 h. Cells were briefly permeabilized with 0.005% Triton X-100, rinsed with PBS, and incubated for 30 min with Click-iT EdU Alexa Fluor 594 Imaging Kit according to manufacturer protocol (Thermofisher, Inc). Stained coverslips were washed once with PBS, counterstained with Hoechst (Life Technologies), mounted in Vectashield mounting media (Vector Laboratories) and imaged by fluorescence microscopy.

### RNA isolation, library preparation and RNAseq analysis

Cell pellets were resuspended in 1 ml RNAlater and placed at 4°C ON. RNA was prepared using the Qiagen RNeasy kit with additional on column DNAase digestion. 10 ug of total RNA was converted into cDNA using an Illumina RNA-seq Prep Kit (Illumina). The experiment was performed in duplicate, resulting in 18 cDNA libraries. cDNA libraries spanning 1.5 cell cycles, and bar-coded samples subjected to high throughput sequencing on an Illumina-HiSeq 2500v4 platform (35M paired end reads/library, two biological replicates per timepoint). Adapters and low quality reads were removed by Trimmomatic (Bolger, Lohse et al. 2014), paired reads with concordant mappings were aligned to the *T. thermophila* macronuclear genome (SB210, 2021 version, TGD) by HISAT2, transcript abundance in each sample was computed using StringTie (Pertea, Kim et al. 2016). Quantification and statistical inference of changes between timepoints were computed by DESeq2 (Love, Huber et al. 2014). Plotting was based on regularized log (rlog) transformed DESeq2 data.

### Identification of cell cycle regulated genes

To identify cell cycle regulated genes, Metacycle was used (Wu, Anafi et al. 2016). MetaCycle implements JTK_CYCLE (JTK) and Lomb-Scargle (LS) and integrates their results. First, genes with two 1.5-fold change in opposite direction between two timepoints was filtered out (p-value < 0.05) using the logarithmic fold changes results by DESeq2. 3825 filtered genes with normalized counts from DESeq2 with biological duplicates were identified and inputted into MetaCycle. MetaCycle::meta2d with minimum and maximum period lengths set as 120 and 240 min, respectively. 3032 genes were detected with FDR<0.05. The 0 min timepoint was excluded from the MetaCycle analysis due to the transient effect of elutriation on the expression of many metabolic genes. LS and JTK algorithms’ performance on peaked data was much lower (Deckard, Anafi et al. 2013).

### Cluster analysis

Cluster analysis was performed with noise-robust soft clustering (Futschik and Carlisle 2005). Soft clustering is more noise robust and generates accessible internal cluster structures. It was implemented using the fuzzy c-means algorithm. Regularized log (rlog) transformed data from DESeq2 was used as input. Optimized FCM parameter m= 1.37114 was used.

### Analysis of gene functions

Gene Ontology (GO) over-representation test of gene clusters was done using PANTHER (Protein Analysis Through Evolutionary Relationships) which contains the complete sets of protein coding genes and reports orthologs and paralogs (GO Ontology database Released 2016-08-22) (Mi, Poudel et al. 2016). Clustering results were inputted into PANTHER. Dotplot comparing pathways of clusters was plotted by ggplot2 in R. Duplicate biological processes having the same genes and similar process names were removed when generating graphs. Additional gene annotation and ontology information was found using TGD (Tetrahymena Genome Database; www.ciliate.org).

### Co-expression profiling

The R package “WGCNA” (Zhang and Horvath, 2005) was used to perform weighted gene co-expression analysis (WGCNA). RNA-seq data were properly pre-processed using Deseq2 to generate vsd values for each sample as recommended. Before construction of the adjacency matrix, a soft threshold (β) was set by inspection of plots generated after calling the function pickSoftThreshold. The soft threshold was set to 6, where the Scale-Free Topology (SFT) Index as a function of the Soft Threshold reached saturation. Modules were generated after calling the function blockwiseModules. Arguments of this function were kept as default. Lists of top 10 co-expressed genes were generated for all cell cycle-regulated genes.

## DATA ACCESS

All raw and processed sequencing data generated in this study have been submitted to the NCBI Gene Expression Omnibus (GEO; https://www.ncbi.nlm.nih.gov/geo/) under accession number GSE123456. All relevant data are within this paper and its Supporting Information files.

## ACKNOWLEDGEMENTS

This work was supported by NSF grant MCB-0132675 and the Tom and Jean McMullin Endowed Professorship to GMK.

---

**Supplemental Figure S1.**
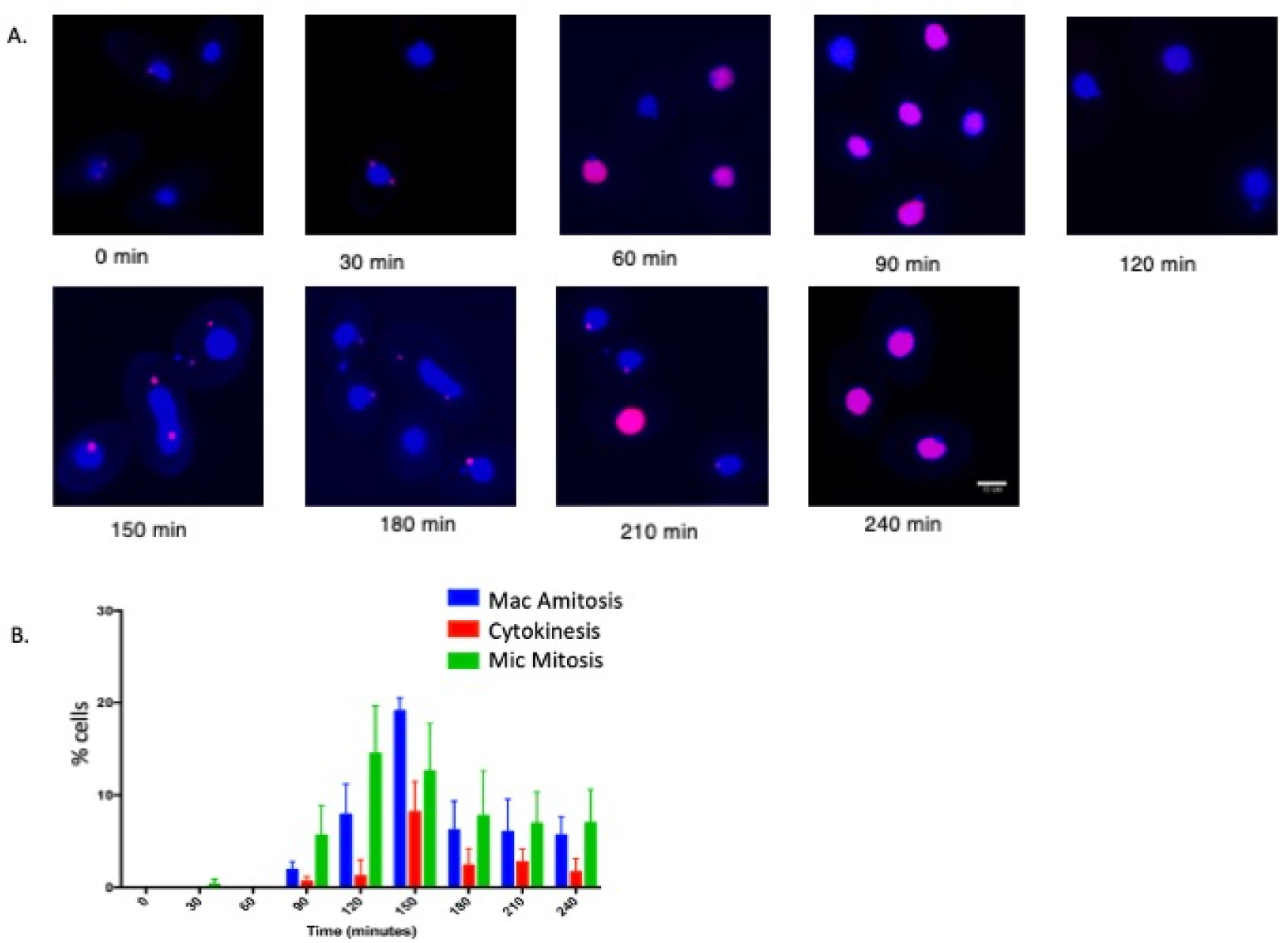
Imaging of DNA synthesis, nuclear division and cell division in elutriated cell populations. **A.** Cells isolated at 30 min intervals after elutriation were pulse labeled with EdU for 15 min and incorporation was visualized microscopically after treatment with fluorescence “click” chemistry reagents. **B.** Percentage of cells undergoing micro- or macronuclear division or cytokinesis. Cell populations isolated at 30 min intervals following elutriation were examined with DAPI under fluorescence microscopy to calculate the percentage of cells in micronuclear mitosis (green bar), macronucleus amitosis (blue bar) or cytokinesis (red bar).

**Supplemental Figure S2.**
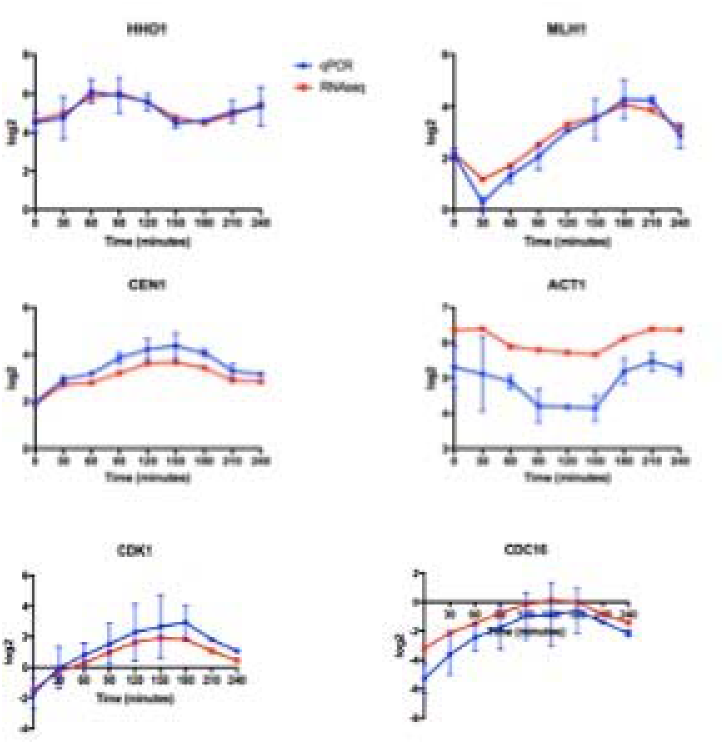
qRT-PCR validation of six cell cycle regulated genes identified by RNA-seq. Graphic representation of RNA-seq and Q-RT-PCR analysis of 6 genes cell cycle regulated genes. For RNA-seq, HHO1, MLH1, CDK1 and CDC16 displayed >2-fold changes in mRNA abundance. CEN1 and ACT1 displayed >1.5-fold changes.

**Supplemental Figure S3.**
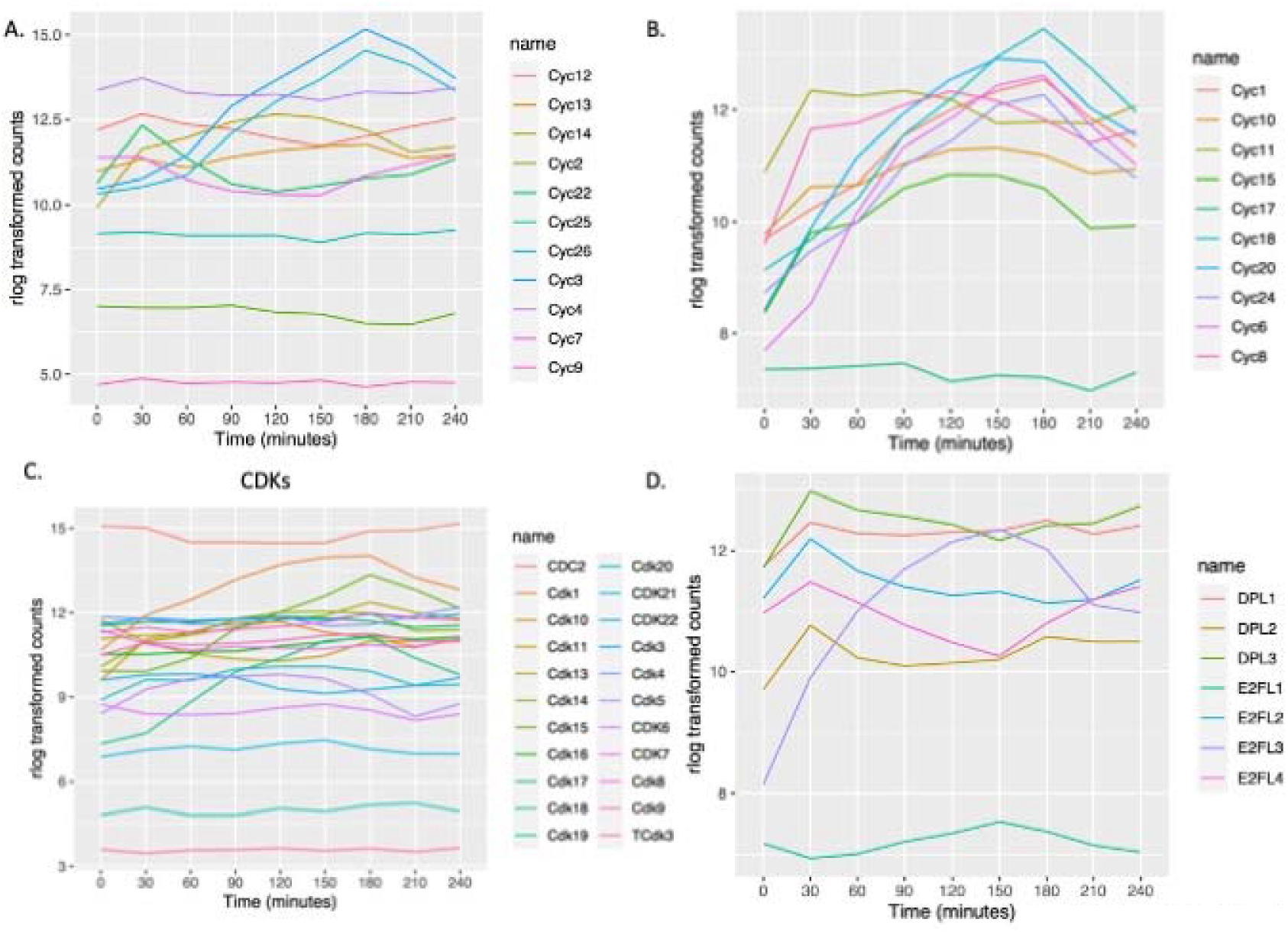
Gene expression profile of cyclin and CDK and E2F genes. **A.** Expression profile of all predicted cyclin D family genes. **B**. Expression profile of all predicted cyclin A/B family genes. **C.** Expression profiles for a predicted CDK genes. **D.** Expression profiles for all E2F family members.

**Supplemental Figure S4.**
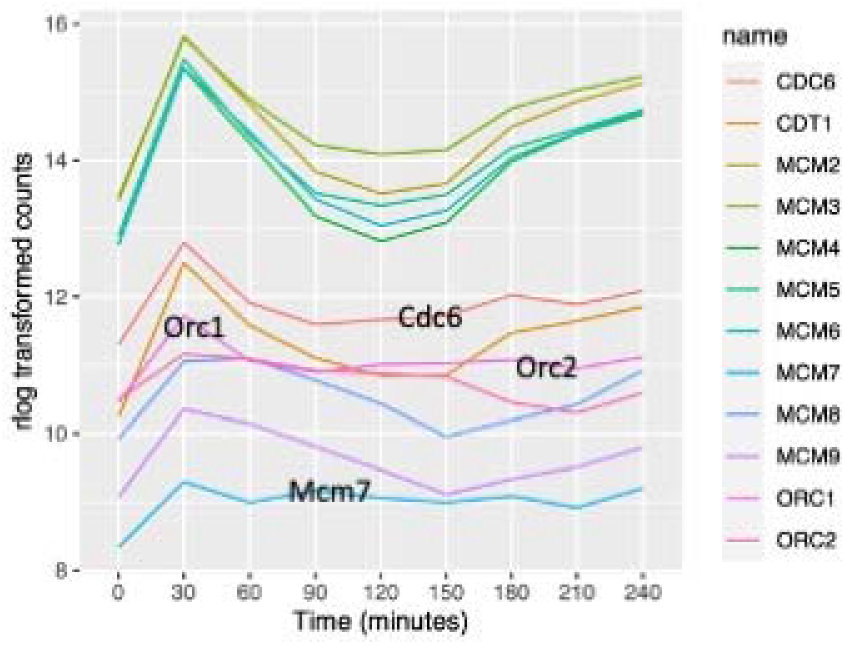
Expression profile of all cell cycle regulated pre-RC components.

